# *In silico* assessment of 18S rDNA metabarcoding markers for the characterization of nematode communities

**DOI:** 10.1101/2023.08.01.551472

**Authors:** Gentile Francesco Ficetola, Alessia Guerrieri, Isabel Cantera, Aurelie Bonin

## Abstract

Nematodes are keystone actors of soil, freshwater and marine ecosystems, but the complexity of morphological identification has limited broad-scale monitoring of their biodiversity. DNA metabarcoding is increasingly used to assess nematode biodiversity but requires universal primers with high taxonomic coverage and high taxonomic resolution. Several primers have been proposed for the metabarcoding of nematode diversity, many of which target the 18S rRNA gene. *In-silico* analyses have a great potential to assess key parameters of primers, including their taxonomic coverage, resolution and specificity. Based on a recently-available reference database, we tested *in-silico* the performance of fourteen commonly used and one newly optimized primer for nematode metabarcoding. Most primers showed very good coverage, as amplified most of sequences in the reference database, while four markers showed limited coverage. All primers showed good taxonomic resolution. Resolution was particularly good if the aim was the identification of higher-level taxa, such as genera or families. Overall, species-level resolution was higher for primers amplifying long fragments. None of the primers was highly specific for nematodes as, despite some variation, they all amplified a large number of other eukaryotes. Differences in performance across primers highlight the complexity of the choice of markers appropriate for the metabarcoding of nematodes, which depends on a trade-off between taxonomic resolution and the length of amplified fragments. Our *in-silico* analyses provide new insights for the identification of most appropriate primers, depending on the study goals and the origin of DNA samples. This represents an essential step to design and optimize metabarcoding studies assessing nematode diversity.

## INTRODUCTION

Nematodes are probably the most abundant animals on Earth, and are a crucial component of soil, freshwater and marine ecosystems (Crowther et al. 2019, van den Hoogen et al. 2019, Delgado-Baquerizo et al. 2020). Despite their importance, the complexity and labor of morphological identification has long limited broad-scale analyses of nematode biodiversity (Porazinska et al. 2009, Sapkota and Nicolaisen 2015). The biodiversity of nematodes is estimated to be 1-10 million species, but less than 30,000 of them have been described using morphology (Hodda 2022a). Recent advances in DNA metabarcoding have fostered the analysis of nematode biodiversity from a range of environments, highlighting their impressive diversity and the multiple key roles they play (Kerfahi et al. 2016, Ohlmann et al. 2018, Crowther et al. 2019, Delgado-Baquerizo et al. 2020, Kawanobe et al. 2021, Zawierucha et al. 2021, Gattoni et al. 2023).

The identification of appropriate primers is a fundamental step of all DNA metabarcoding analyses (Taberlet et al. 2018, Ficetola and Taberlet 2023). Several features are extremely important for the selection of primers. First, primers must have a low number of mismatches with the sequences of the target group (high taxonomic coverage; Ficetola et al. 2010, Piñol et al. 2015, Marquina et al. 2019). Second, primers should amplify highly variable regions, to enable the identification of target taxa at a high taxonomic level (high resolution; Ficetola et al. 2010, Marquina et al. 2019). Third, traditional metabarcoding analyses generally amplify short genomic regions to reduce the costs associated with sequencing. Even though recent advances in sequencing technologies are leveraging these limitations (Jamy et al. 2022), the use of primers amplifying short fragments is relevant when working with environmental and ancient DNA, as it is often degraded and consists of short sequences (Eichmiller et al. 2016, Jo et al. 2017, Walker et al. 2017, Bylemans et al. 2018, Taberlet et al. 2018). Finally, primers that only amplify the target taxon are often preferred (i.e. primers with high specicitity), because a-specific amplification can reduce the detection of target taxa, particularly if they show limited abundance (Ficetola et al. 2010, Coissac et al. 2012, MacDonald and Sarre 2017, Taberlet et al. 2018).

*In-silico* approaches are extremely useful to assess key features of primers, including the number of mismatches with the target sequences (a key determinant of taxonomic coverage) and the potential taxonomic resolution (Ficetola et al. 2010). *In-silico* tests allow cheap and rapid comparisons of a very large number of primers and often provide a good estimate of the actual performance of primers across various taxa, that can later be confirmed by *in-vitro* assays on real-worlds samples (Ficetola et al. 2010, Epp et al. 2012, Clarke et al. 2014, Collins et al. 2019, Kawanobe et al. 2021, Van Nynatten et al. 2023). Accurate *in-silico* assessments of primers for DNA metabarcoding require the availability of extensive, high-quality reference databases over which primers can be tested (Ficetola et al. 2010, Collins et al. 2019). The recent publication of a curated 18S rRNA database of nematode sequences (18S-NemaBase; Gattoni et al. 2023) poses the basis for such assessments.

In this study, we built upon the 18S-NemaBase to compare the performance of 15 primers proposed for the metabarcoding of nematodes (Table 1). Using *in-silico* PCR, we 1) assessed whether the selected primers are able to amplify a large proportion of nematode taxa (coverage), 2) tested the taxonomic resolution of primers and evaluated whether there is a trade-off between marker length and taxonomic resolution and 3) tested primer specificity, i.e., assessed whether they only amplify nematodes, or also amplify a broad range of other organisms. Our results help to evaluate the appropriateness of different primers for different aims, ranging from the analysis of potentially degraded environmental DNA (eDNA) to whole-organism community DNA (Pawlowski et al. 2020).

**Table 1.**
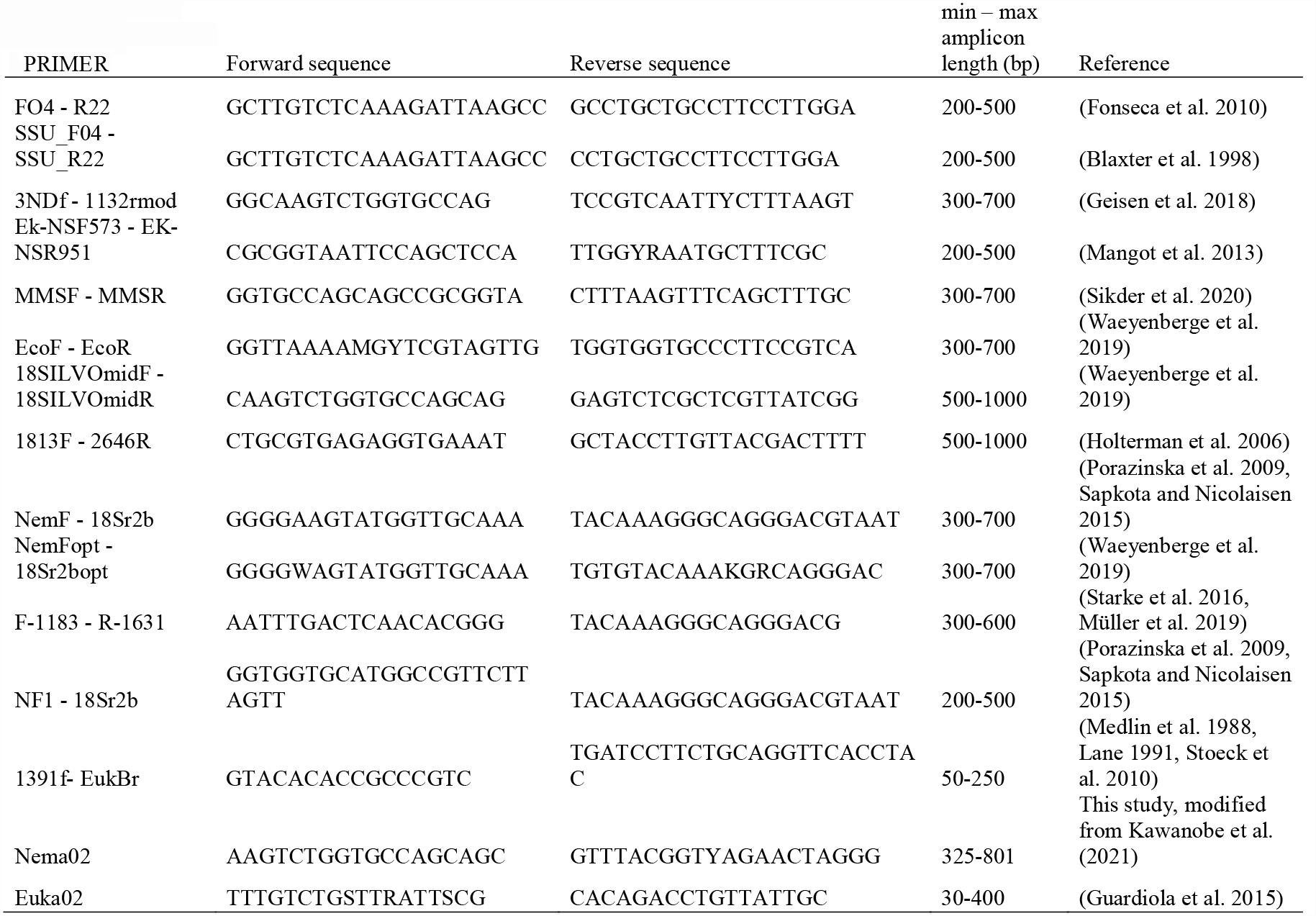
List of primers tested in-silico, forward and reverse sequences of primers and the expected minimum and maximum lengths (bp) of the amplicons used in the ecoPCR program.

## METHODS

### Primer selection

All 15 primers selected for our analyses target the 18S rRNA region of nematodes (Table 1). Among these primers, 13 were selected because they have been identified by a previous review as the primers more commonly used in nematode metabarcoding (Gattoni et al. 2023). One additional primer, Euka02 (Guardiola et al. 2015) is often used for metabarcoding of eukaryotic eDNA and it has been suggested to provide a good estimate of nematode diversity (Guardiola et al. 2015, Ohlmann et al. 2018, Rosero et al. 2021, Calderón-Sanou et al. 2022). Furthermore, we developed a new primer pair (Nema02; see Table 1) by optimizing primer pair F548_A / R1912 from Kawanobe et al. (2021). More specifically, we performed in silico PCRs on the public sequence database GenBank v249 with the ecoPCR program (Ficetola et al. 2010) to evaluate Nematoda and non-Nematoda variability at each position of the sequences matching the F548_A and R1912 primers, and in a 10-base interval in 5’ and 3’ of these sequences. The objective was to fine-tune the primer sequences to maximize non-Nematoda variability while minimizing Nematoda variability in primer-matching sequences, especially in 3’, in order to increase specificity. Taxonomic resolution of the associated marker was evaluated using the ecostaxspecificity program of the OBITools package (Boyer et al. 2016). Compatibility of annealing temperatures and absence of problematic primer dimers or hairpins were checked using OligoCalc (http://biotools.nubic.northwestern.edu/OligoCalc.html) (Kibbe 2007) and the OligoAnalyzer Tool (https://eu.idtdna.com/pages/tools/oligoanalyzer?returnurl=%2Fcalc%2Fanalyzer), respectively.

### In-silico PCR

The availability of high-quality, curated databases is pivotal to test the performance of metabarcoding primers. We based our analyses on the 18S-NemaBase (Gattoni et al. 2023), which represents the most complete and high-quality available reference database of nematode 18S rRNA. The original 18S-NemaBase includes 5 231 sequences identified at family level or better, representing 214 families and 2734 species (Gattoni et al. 2023).

The performance of primers was tested using the ecoPCR program (Ficetola et al. 2010). EcoPCR allows *in-silico* assessment of amplification of a sequence on the basis of its match with a selected primer pair in a region of a specified length (Ficetola et al. 2010). The sequence is selected from a given reference database. To work, the program requires the reference database in the ecoPCR format. Thus, we converted the 18S-NemaBase from the fasta to the ecoPCR format using the obiconvert command of the OBITools command suit (Boyer et al. 2016). The original 18S-NemaBase database consisted of 5231 sequences.

However, 283 sequences (i.e. 5,4%) were excluded because they showed problems during the conversion from the fasta to the ecoPCR format (268 sequences) or because they were they were assigned to non-nematode taxa in 18S-NemaBase (15 sequences), thus we based our analyses on a total of 4948 sequences.

For *in-silico* PCR, we allowed a maximum of 3 mismatches between each primer and the sequences in the database. For each primer pair, the length of amplified fragments was selected based on the literature and of preliminary trials (Table 1).

Running ecoPCR on the curated 18S-NemaBase allowed us to calculate two measures of primer performance:

- Taxonomic coverage, as the percentage of amplified sequences within the reference database
- Resolution, i.e. the ability of markers to distinguish between closely-related taxa

Taxonomic resolution was calculated using the procedure detailed in Ficetola et al. (2021). First, all the sequences obtained in each EcoPCR were compared among them to produce a list of unique metabarcodes. We then obtained the list of taxa associated to each unique metabarcodes. Taxonomic resolution was tested at three levels: species, genus and family. We assessed if, for each unique metabarcodes and taxonomic level, all the amplified taxa belong to the same taxon. Let’s assume, for instance, that multiple species within one genus share the same metabarcode. This particular metabarcode shows a genus-level and a family-level resolution, but not a species-level resolution. The average taxonomic resolution of markers was then calculated as the proportion of unique metabarcodes that have a species-level, genus-level and family-level resolution (Ficetola et al. 2021).

We used a linear regression to test whether there is a positive relationship between taxonomic resolution and the log-transformed average length of the amplified fragments. This analysis was run at the species-level resolution because all the primes showed excellent resolution at the genus- and family level (see results).

Furthermore, we assessed the specificity of the tested primers using the whole GenBank database (version 249) instead of the 18S-NemaBase as reference database in the ecoPCR program. This time, we only retained one sequence per species, to avoid biased due to the overrepresentation of model species (e.g. *Caenorhabditis elegans* for nematodes). Then, specificity was measured as the percentage of nematodes amplified over non target organisms (Ficetola et al. 2010). Primers showing high specificity are in principle preferable because they have a higher probability of detecting target taxa, including rare species (Ficetola et al. 2010, Wilcox et al. 2013, Taberlet et al. 2018, Collins et al. 2019, Leese et al. 2021).

## RESULTS

### Taxonomic coverage

The taxonomic coverage (proportion of amplified sequences compared to the 18S-NemaBase database) was highly variable across primers. The majority of primers showed very good to excellent coverage, amplifying >90% of sequences in the reference database, while four primers showed a limited coverage (Table 2, Fig. 1). The three primers with the highest coverage were 3NDf-1132rmod, EcoF-EcoR and Euka02, all showing coverage ≥ 97%. These primers amplified fragments with very different lengths, spanning from 100 bp (Euka02) to more than 800 bp (1813F_2646R; Table 2).

**Table 2.**
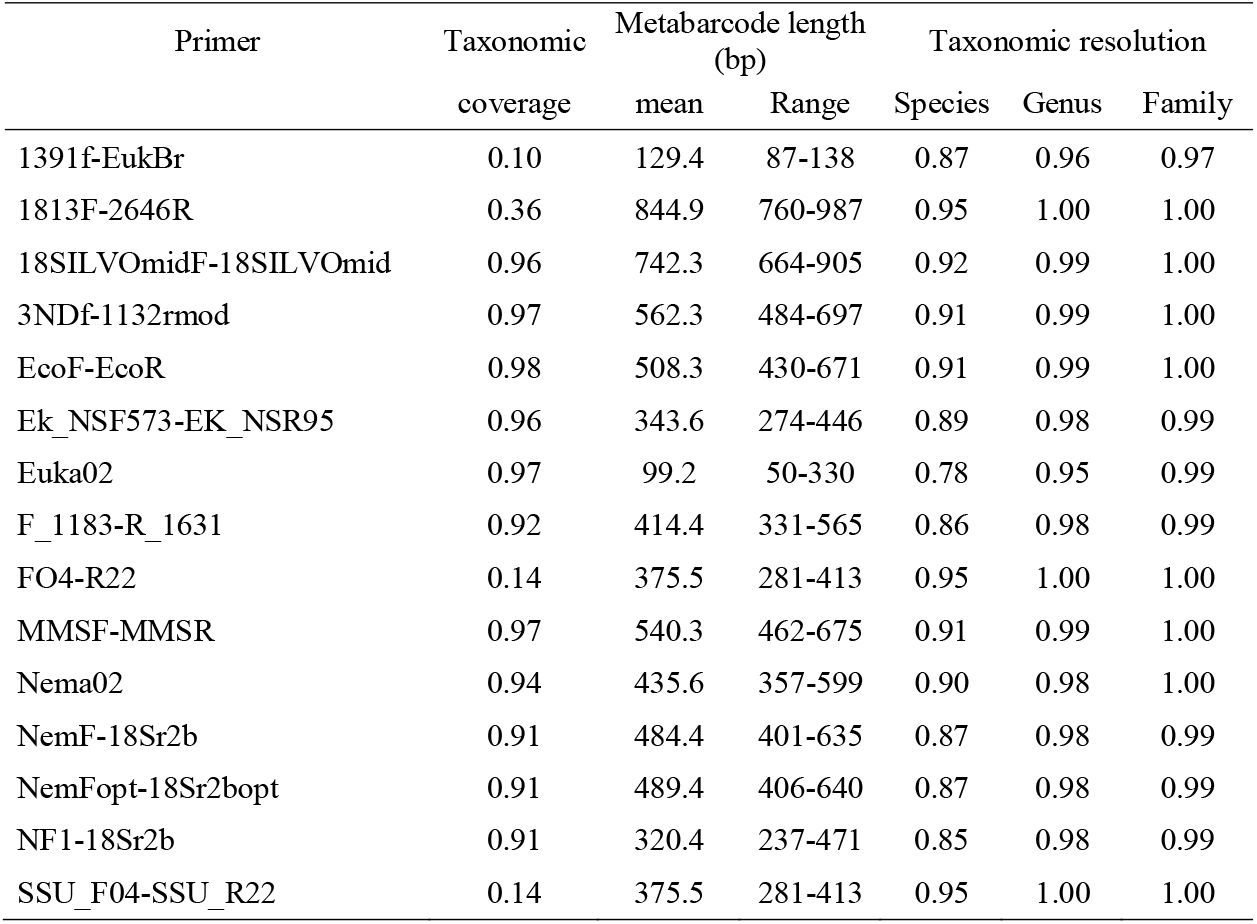
Results of *in-silico* PCRs testing the taxonomic coverage and the resolution of 15 primers proposed for nematode metabarcoding.

**Figure 1.**
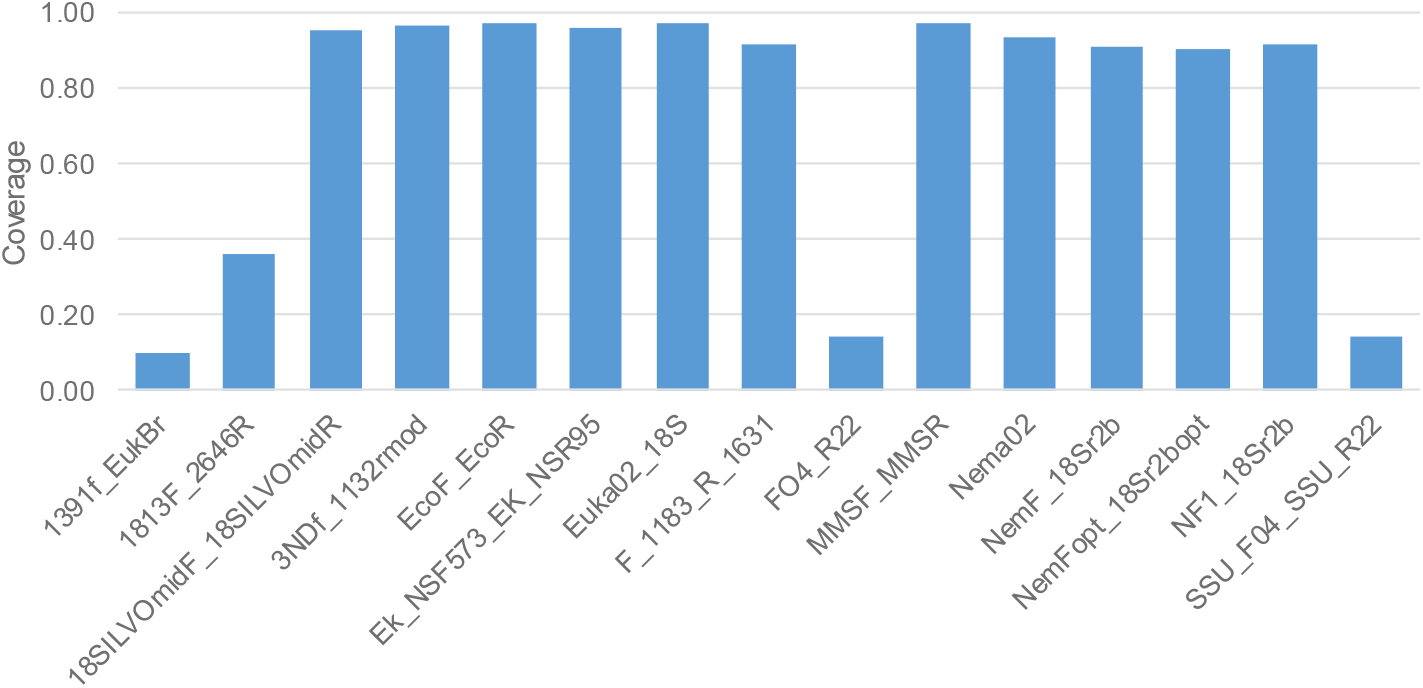
Taxonomic coverage of the 15 primers tested *in-silico* on the 18S-NemaBase.

### Taxonomic resolution

All primers showed a good resolution if the aim was species-level identification. Even the primer with the lowest species-level resolution (Euka02), associated most (78%) of unique metabarcodes with just one nematode species (Table 2, Fig. 2). Eight primers showed a species-level resolution >90%; all of them amplified fragments >350bp. The three primers with the highest species-level resolution (95%) showed generally low coverage (Table 2, Fig. 3).

**Figure 2.**
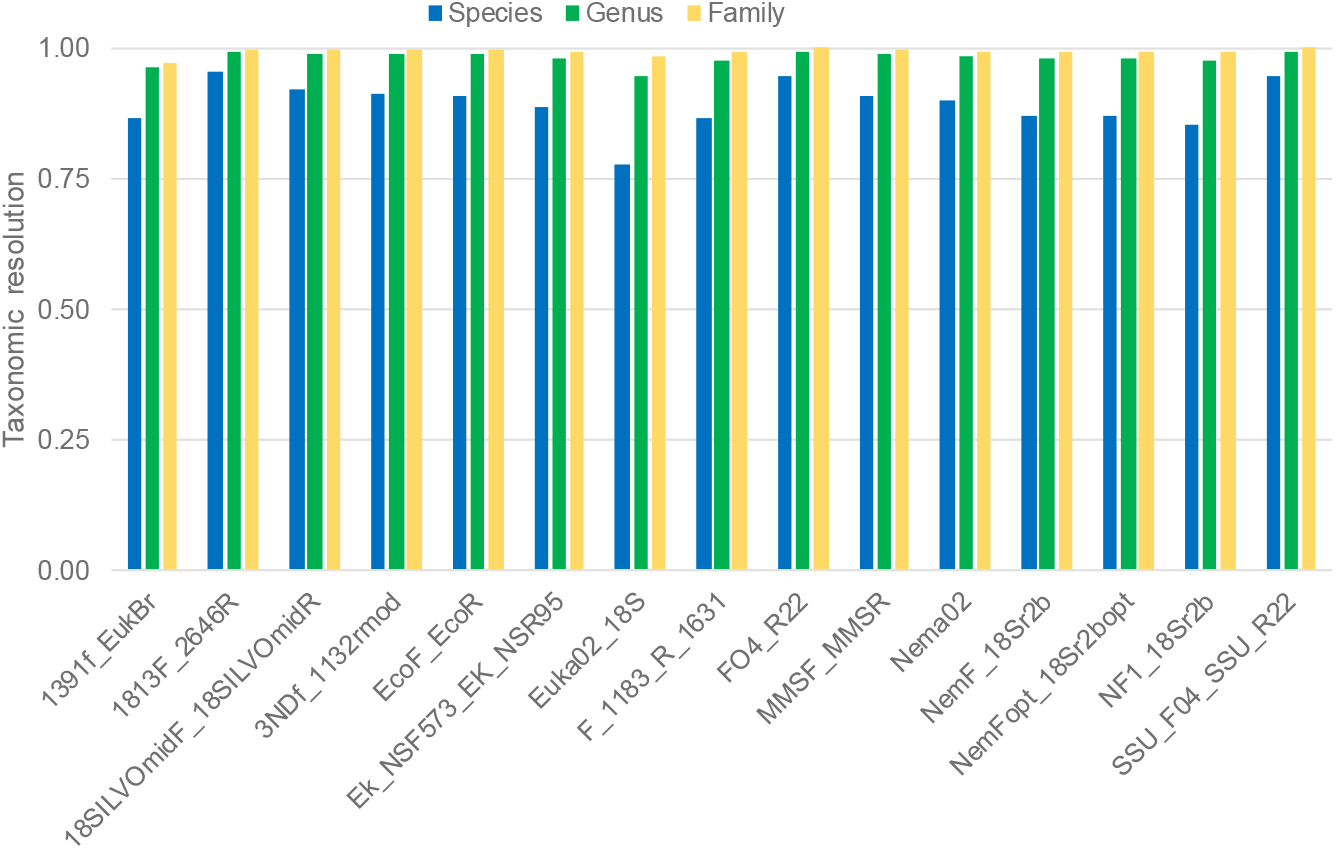
Species-, genus- and family- level taxonomic resolution of the 15 primers tested *in-silico* on the 18S-NemaBase.

**Figure 3.**
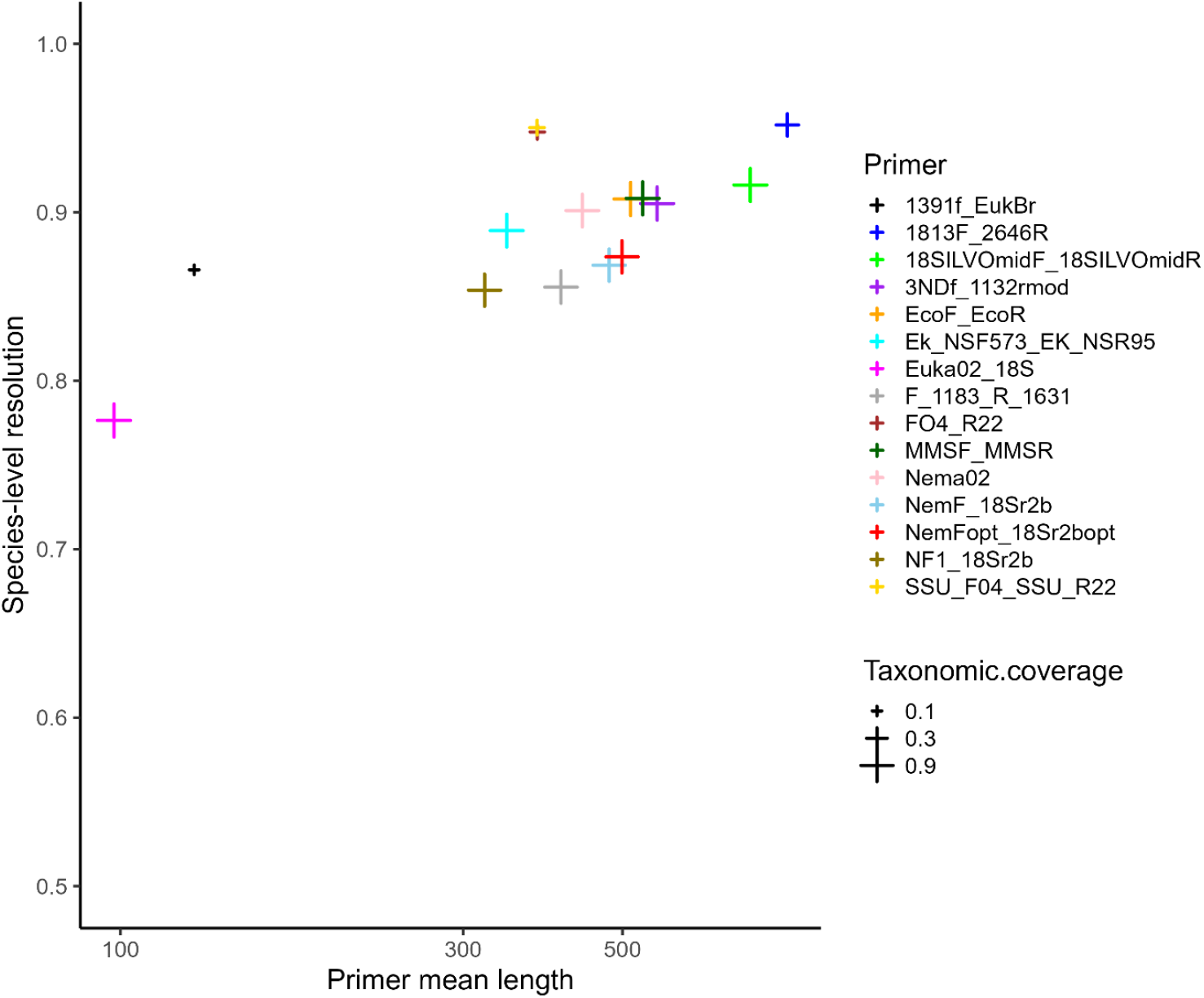
Relationship between mean length (bp) of amplified fragments and species-level resolution of the 15 primers. The size of symbols is proportional to primers coverage

If the aim was genus-level identification, all primers showed excellent resolution, as they were able to tease apart 95% of genera or more. Resolution was even better if the target was family identification, with all primers showing a resolution ≥97% (Table 2, Fig. 2).

Overall, we observed a positive relationship between primer taxonomic resolution at the species level, and the length of the amplified metabarcodes (linear regression: *F*_1,13_ = 11.92, *P* = 0.004; *R*^2^ = 0.48; Fig. 3), suggesting that the increase in the metabarcodes length facilitate taxonomic resolution at the species-level.

### Specificity

When tested on the whole GenBank, primers showed specificity values that ranged between 1% or less (FO4-R22, SSU_F04-SSU_R22) and ∼7% (1813F-2646R, Nema02; Fig. 4a).

**Figure 4.**
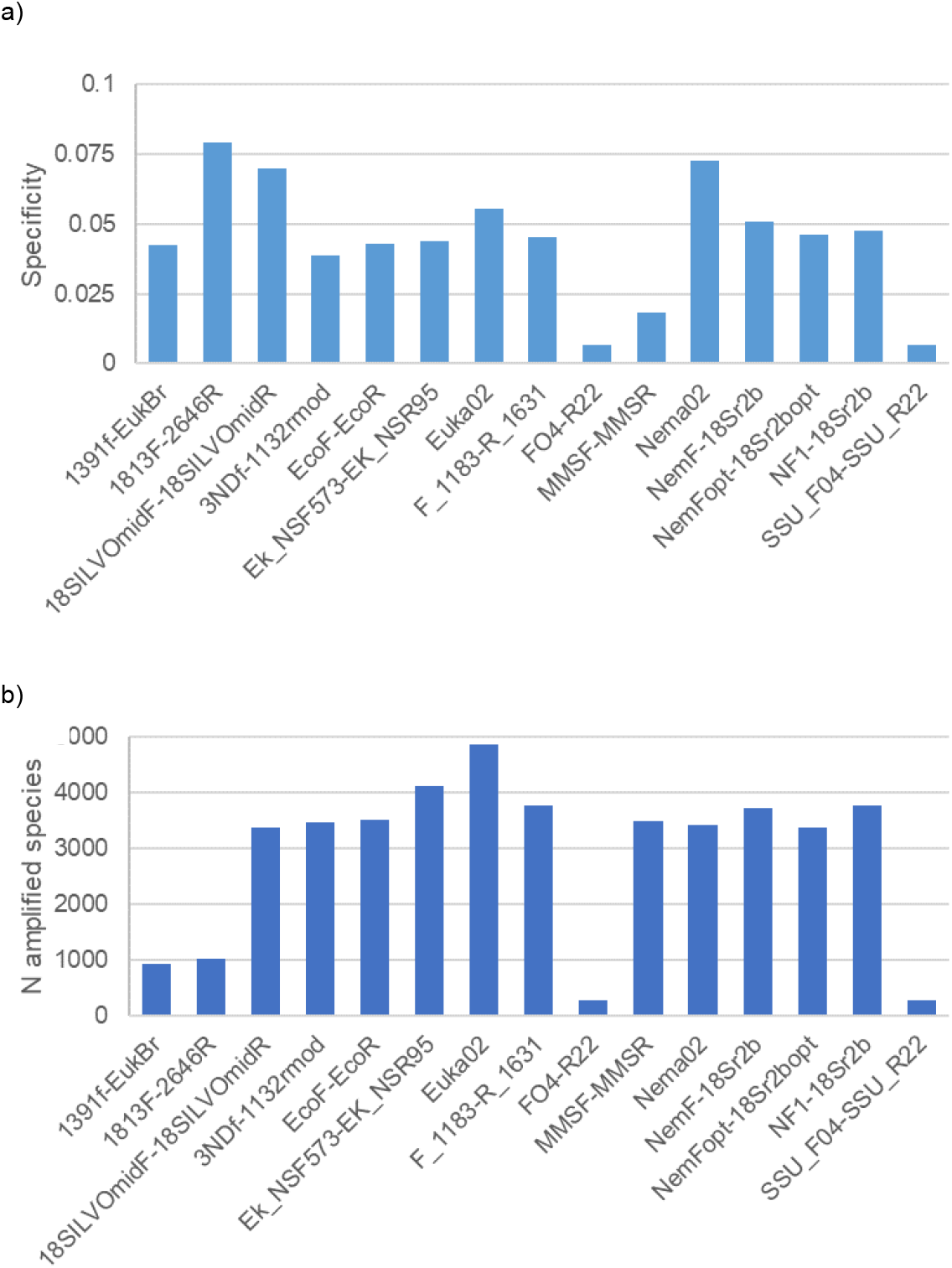
a) Specificity of the 15 primers tested *in-silico* on the whole Genbank. Specificity was measured as the N of nematode sequences / total N of amplified sequences. b) Number of nematode species amplified by the different primers, when the *in-silico* PCR is run on the whole GenBank

These values indicate that all the markers amplify a very large number of non-nematode sequences. Nevertheless, some of the primers with high specificity only amplified a limited number of nematode taxa (Fig. 1, Fig. 4b). The primers with the best compromise between specificity and coverage of nematodes included 18SILVOmidF-18SILVOmid, Euka02 and Nema02 (Fig. 2, Fig. 4).

## DISCUSSION

Our analyses showed that primers suggested for the analysis of nematode diversity have heterogeneous performances, particularly in terms of taxonomic coverage and specificity. Conversely, the taxonomic resolution of all primers was generally good, and across-primer variation in resolution was strongly related to the well-known trade-off with metabarcode length (Fig. 3). The selection of the most appropriate primer pair for the analyses of nematode biodiversity depends on the balance between multiple factors, including study aims, media from which DNA is extracted, and adopted sequencing technology.

The taxonomic coverage was highly variable. The majority of primers showed very good to excellent coverage, as they amplified more than 90% (and sometimes >97%) of available sequences, still, some showed limited coverage. The primers with low coverage were either universal primers that amplify large bunch of eukaryotes and are not designed specifically for nematodes (Medlin et al. 1988, Lane 1991, Fonseca et al. 2010), or primers from landmark studies of nematode phylogeny (Blaxter et al. 1998, Holterman et al. 2006), that were developed using sequences available at that time. It should be highlighted that our bioinformatics analyses are based on the number of mismatches in the priming region. This parameter is particularly relevant in metabarcoding studies, when DNA is amplified by complex mixtures comprising the DNA of many different taxa. In these cases, a large number of mismatches is problematic, as species with less mismatches are amplified preferentially, while species with less mismatches tend to be overlooked (Piñol et al. 2015, Fonseca 2018, Marquina et al. 2019). A few mismatches probably are less problematic in phylogenetic studies, where the DNA of just one species is extracted at each time directly from specimens (Blaxter et al. 1998, Holterman et al. 2006).

*In-silico* assessments of primer coverage, like the one performed here, have their own limitations, as they may both miss some taxa that would be amplified *in-vitro* and vice versa. *In-silico* analyses focusing on curated databases can also miss key issues, such as non-specific amplification (Collins et al. 2019). Nevertheless, several *in-vitro* tests have confirmed the appropriateness of primers studied with *in-silico* analyses (Epp et al. 2012, Clarke et al. 2014, Kartzinel and Pringle 2015, Collins et al. 2019, Waeyenberge et al. 2019, Kawanobe et al. 2021). For instance, our results are in agreement with Sikder et al. (2020), which used mock communities to compare the performance of MMSF-MMSR, NemF-18Sr2b and SSU_F04-SSU_R22 and observed a limited coverage of SSU_F04-SSU_R22. Similarly, Kenmotsu et al. (2021) confirmed that F_1183-R_1631 and NF1_18Sr2b show a similar, very good performance; Geisen et al. (2018) confirmed the excellent performance of 3NDf-1132rmod; and Guardiola et al. (2015) suggested an excellent coverage for Euka02 (see also Ficetola et al. 2021 for in-vitro tests confirming the excellent taxonomic coverage of this marker for all the tested invertebrate phyla). Nevertheless, comparative *in-vitro* tests performed on both mock communities and real samples will be extremely important to validate our conclusions on taxonomic coverage, particularly for primers that have received limited testing so far (e.g., Nema02, but se Kawanobe et al. 2021 for analyses confirming the good performance of similar primers).

All primers showed good to excellent resolution on the considered reference database. Even the shortest primer (Euka02) showed a reasonably good ability to discriminate between species (Fig. 2) and, for all primers, the frequency of genera sharing the same metabarcode was ≤5% (i.e. genus-level resolution was always 95% or higher). These findings are highly promising for the use of metabarcoding for nematode analyses. However, it is important to acknowledge some caveats. The use of high quality, curated databases is fundamental for all the metabarcoding analyses, being pivotal for assessments of marker performance, and for accurate taxonomic identification (Taberlet et al. 2018, Leray et al. 2019, Morinière et al. 2019, White et al. 2020, Williams et al. 2023). Our analyses were run on a large, curated database containing about 5000 sequences from 214 families, 668 genera and 2734 species (Gattoni et al. 2023). Unfortunately, this database only represents the currently described species and genera, but most nematodes inhabiting the Earth still require description (Hodda 2022a, Hodda 2022b). We stress that all measures of taxonomic resolution strongly depend on the available data (Weigand et al. 2019). For instance, if the reference database only includes one species within a given genus, analyses would return a species-level resolution, despite unanalysed species within that genus may share the same metabarcodes (Ficetola et al. 2021).

Within the primers with good taxonomic coverage, resolution was clearly related to the length of the amplified fragment (Fig. 3). This is not unexpected, as longer fragments include more variable sites and thus are well known to provide a better resolution (Taberlet et al. 2018). Nevertheless, even the shortest fragments allow an excellent genus-level resolution. Many functional analyses of nematodes are performed at the genus level. For instance, Nemaplex (http://nemaplex.ucdavis.edu) is a major database of nematode traits, and provides traits at the genus-level resolution. Ensuring that primers provide robust identification at the genus level is thus extremely important for all the studies focusing on nematode traits and functional diversity.

A key issue of the analyzed primers is that they are not specific to nematodes, and amplify a broad range of eukaryotes (Fig. 4). A-specific amplification can reduce the detection of rare taxa, and can even increase false positives (Wilcox et al. 2013, Collins et al. 2019). This can be particularly problematic when analyses target complex mixtures of DNA(e.g. eDNA extracted from soil) that comprise the DNA of both nematodes and other organisms (Jurburg et al. 2021, Geisen et al. 2023). For instance, in-vitro assessments of the EcoF-EcoR primers detected a very large number of non-nematode taxa, suggesting that this marker can be not appropriate for analyses only focusing on nematodes (Waeyenberge et al. 2019). The low specificity of most of primers make them valuable for whole analyses of eukaryote diversity in soil, sediments or aquatic environments (Fonseca et al. 2010, Guardiola et al. 2015, Waeyenberge et al. 2019). Studies focusing on nematodes should thus assess whether the retrieved data are enough to obtain reliable estimates of species diversity / occurrence. Approaches such as rarefaction curves and analyses of detection probability can allow to assess whether key parameters, such as the number of replicates and sequencing depth, are enough to obtain robust biodiversity estimates, or need to be increased for well-grounded ecological inference (Ficetola et al. 2015, Zinger et al. 2019, Jurburg et al. 2021).

Given the trade-off between taxonomic resolution and fragment length, our analysis can help to identify the most appropriate primer pair depending on the study focus. Usually, studies performed on DNA extracted from difficult substrates (e.g. environmental DNA from sediments and water; Taberlet et al. 2018) target short DNA fragments. In fact, DNA from environmental samples is often degraded, and the analysis of short fragments (ideally <150 bp) ensure a better probability of detecting target taxa (Parducci et al. 2017). Short fragments have the advantages of being sequenced with platforms such as Illumina HiSeq and NovaSeq, that produce an impressive number of reads at steadily decreasing prices, and thus enable processing a very large number of samples. Longer fragments (within 450-500 bp) can be appropriate for both metabarcoding of whole-organism community DNA, for non-extracellular eDNA, and for eDNA extracted from substrates protecting eDNA from degradation (e.g. developed soils with high clay content) (Levy-Booth et al. 2007, Pietramellara et al. 2009). These fragments can be sequenced with platforms such as Illumina MiSeq, which still enable processing a large number of samples at reasonable prices. Primers well suited for this aim include 3NDf-1132rmod, MMSF-MMSR and, if confirmed by *in-vitro* tests, Nema02. Among these primers, Nema02 is the one with the highest specificity for nematodes (Fig. 4a-b). Finally, longer fragments will be particularly suited for whole-organism community DNA (but see Jamy et al. 2022 for a remarkable example of long-read analysis of DNA extracted from environmental samples). The continuing developments of high-throughput sequencing are making long-read metabarcoding increasingly feasible at progressively more affordable prices.

Ongoing development of DNA metabarcoding are opening new avenues to the study of biodiversity, and to the identification of management priorities (Eisenhauer et al. 2021, Guerra et al. 2021, Zeiss et al. 2022). Nematodes are increasingly recognized as key components of soil communities, and advances in molecular approaches to assess and monitor their biodiversity are extremely important for the growing knowledge on this highly diverse phylum (Kawanobe et al. 2021, Hodda 2022b). The selection of most appropriate markers is a fundamental step to maximise the information drawn by each study (Zinger et al. 2019).

## Aknowledgments

This study was funded by the European Research Council under the European Community’s Horizon 2020 Programme, Grant Agreement no. 772284 (IceCommunities).

